# Macrophages and glia are the dominant P2X7-expressing cell types in the gut nervous system – no evidence for a role of neuronal P2X7 receptors in colitis

**DOI:** 10.1101/2022.02.02.478704

**Authors:** Tina Jooss, Jiong Zhang, Tanja Rezzonico-Jost, Björn Rissiek, Friedrich Koch-Nolte, Tim Magnus, Susanna Zierler, Michael Schemann, Fabio Grassi, Annette Nicke

## Abstract

**Summary:** Blockade or deletion of the pro-inflammatory P2X7 receptor channel has been shown to reduce tissue damage and symptoms in models of inflammatory bowel disease (IBD) and P2X7 receptors on enteric neurons were suggested to mediate neuronal death and associated motility changes. Here we used P2X7-specific antibodies and nanobodies as well as a BAC transgenic P2X7-EGFP reporter mouse model and *P2rx7^−/−^* controls to perform a detailed analysis of cell type-specific P2X7 expression and possible overexpression effects in the enteric nervous system. In contrast to previous studies, we did not detect P2X7 in neurons but found dominant expression in glia and macrophages which closely interact with the neurons. P2X7 overexpression *per se* did not induce significant pathological effects. Our data indicate that macrophages and/or glia account for P2X7-mediated neuronal damage in IBD and provide a refined basis for the exploration of P2X7-based therapeutic strategies.

## Introduction

Inflammatory bowel diseases (IBD), such as ulcerative colitis and Crohn’s disease, are chronic relapsing-remitting inflammatory diseases with rising incidence in industrialized countries. They are thought to result from an abnormal host-microbial response of the intestinal immune system in a genetically susceptible host (Baumgart and Carding, 2007; Orholm et al., 1991). The resulting imbalance of homeostatic immunomodulatory mechanisms is characterized by increased macrophage and T-cell activation and up-regulation of proinflammatory cytokines (Sanchez-Munoz et al., 2008; Xavier and Podolsky, 2007). In addition, IBDs are associated with enteric neurodegeneration and abnormal gut motility and the resulting breakdown of enteric nervous system (ENS) control is a major contributing factor in the development of functional bowel disorders (Gulbransen et al., 2012; Niesler et al., 2021; Spear and Mawe, 2019).

Extracellular purines play important roles in immune cell regulation and enteric signaling (Antonioli et al., 2015; Burnstock, 2016; Diezmos et al., 2016; Gulbransen and Sharkey, 2009; Vuerich et al., 2020). High concentrations of ATP are sensed as a danger signal by the P2X7 receptor and it is generally accepted that activation of this non-specific cation channel acts as a secondary stimulus to induce NLRP3 inflammasome assembly and subsequent maturation and release of proinflammatory IL-1β from macrophages. In addition, it has been shown that ATP, via the P2X7 receptor, induces apoptosis in several cell types (Di Virgilio et al., 2017; Di Virgilio et al., 2018; Proietti et al., 2014; Rissiek et al., 2015). While pathophysiological roles of P2X7 receptors have been well established in *P2rx7^−/−^* mouse models (Kaczmarek-Hajek et al., 2012; Rumney and Gorecki, 2020), possible physiological roles are comparably poorly investigated.

P2X7 expression was shown to be increased in IBD patients and animal models, and its deletion or blockade reduced tissue damage in different rodent colitis models (Antonioli et al., 2019; Evangelinellis et al., 2021; Vuerich *et al*., 2020). In a short phase IIa clinical study with a limited number of patients, oral application of an P2X7 antagonist ameliorated the pain component in the Crohn’s disease activity index but did not change biomarkers of inflammation (Eser et al., 2015).

A variety of cell types in the colon has been shown to express P2X7 receptors and were involved in colitis pathology. These include immune cells like mast cells (Kurashima et al., 2012; Matsukawa et al., 2016; Ohbori et al., 2017) and different types of T cells (Figliuolo et al., 2017; Schenk et al., 2011), epithelial cells (Cesaro et al., 2010) (Hofman et al., 2015) (Diezmos et al., 2018), and enteric neurons of both the submucosal and myenteric plexus (da Silva et al., 2017; Palombit et al., 2019) (Antonioli et al., 2014; Evangelinellis *et al*., 2021; Gulbransen *et al*., 2012). P2X7-mediated neuronal cell death was implicated in motor disfunction associated with colitis (da Silva *et al*., 2017; Palombit *et al*., 2019) (Antonioli *et al*., 2014; Brown et al., 2016; Gulbransen *et al*., 2012). Accordingly, elevated levels of extracellular ATP chronically activate a neuronal P2X7/Pannexin/Asc/caspase complex that leads to preferential loss of nitrergic neurons (Gulbransen *et al*., 2012). While numerous mechanisms and cell types have been proposed to mediate protective effects of P2X7 blockade, localization of P2X7 receptors in specific cell types in *situ has* been challenging due to a lack of metabolically stable and selective pharmacological tools. In addition, specificity of the used antibodies has not been demonstrated in most studies. So far, the role of P2X7 receptors in neurons remains unclear but its cell type-specific expression, in particular in neurons versus immune cells, has important implications for potential treatments. In this study, we therefore set out to perform a detailed analysis of P2X7-expressing cell-types in the colon using a BAC transgenic P2X7-EGFP reporter mouse model (Kaczmarek-Hajek et al., 2018) in combination with a P2X7-selective nanobody and *P2rx7^−/−^* mice as negative control.

## Results

### Expression of P2X7 in different layers of the colon

To determine P2X7 localization across the different layers of the colon, we first analyzed cross sections of the distal colon in the P2X7-EGFP transgenic mouse model (Kaczmarek-Hajek *et al*., 2018) (Fig. 1A,B). In this mouse, an EGFP-tagged P2X7 receptor is overexpressed under the control of BAC-derived P2X7 promoters. The respective BAC contains the full-length *P2rx7* gene flanked by about 100 kb and 10 kb up- and downstream sequences, respectively, and careful analysis of P2X7-EGFP localization in the central nervous system (CNS) has previously confirmed an P2X7-EGFP expression pattern that is identical to the endogenous P2X7 receptor and revealed predominant expression in microglia, oligodendrocytes and Bergmann glia (Kaczmarek-Hajek *et al*., 2018). Co-staining of colon slices with antibodies against GFP and P2X7 revealed good co-localization of both signals in single cells throughout the lamina propria (arrow heads in Fig. 1 B) and in two distinct layers adjacent to and in the muscular layer (arrows in Fig. 1 B). The cells in the lamina propria show more intense staining and based on their size, morphology, and localization, represent most likely macrophages, in agreement with the known high P2X7 expression in this cell type (Di Virgilio *et al*., 2018). A weaker EGFP staining and more intense signal of the P2X7 antibody was seen in the apical side of the mucosa. Since the anti-P2X7 antibody should detect both P2X7-EGFP and endogenous P2X7, this could indicate a lower expression of the P2X7-EGFP protein in enterocytes which would be in agreement with the dynamic regulation of P2X7 expression in epithelial cells (Hofman *et al*., 2015). Alternatively, unspecific staining or higher background signal of the P2X7 antibody in these cells or in this preparation could account for the difference. To further confirm epithelial P2X7-EGFP expression, we also performed co-staining of P2X7-EGFP with an anti-P2X7 nanobody in planar mucosa preparations (Fig. 1C). This revealed an almost complete signal overlap, confirming a good correlation between P2X7-EGFP and endogenous P2X7 localization in epithelial cells. As presence of P2X7 receptors in enteric neurons has been frequently reported (Antonioli *et al*., 2014; da Silva *et al*., 2017; Gulbransen *et al*., 2012; Palombit *et al*., 2019) we next performed co-staining for P2X7 and the neuronal markers HuC/D (RNA-binding proteins HuC and HuD) to further determine the identity of the P2X7-expressing cells in the tunica muscularis (Fig. 1D). While both P2X7 and HuC/D signals were present in the same layers, they were clearly separated, indicating localization in different cell types or different subcellular structures.

**Figure 1:**
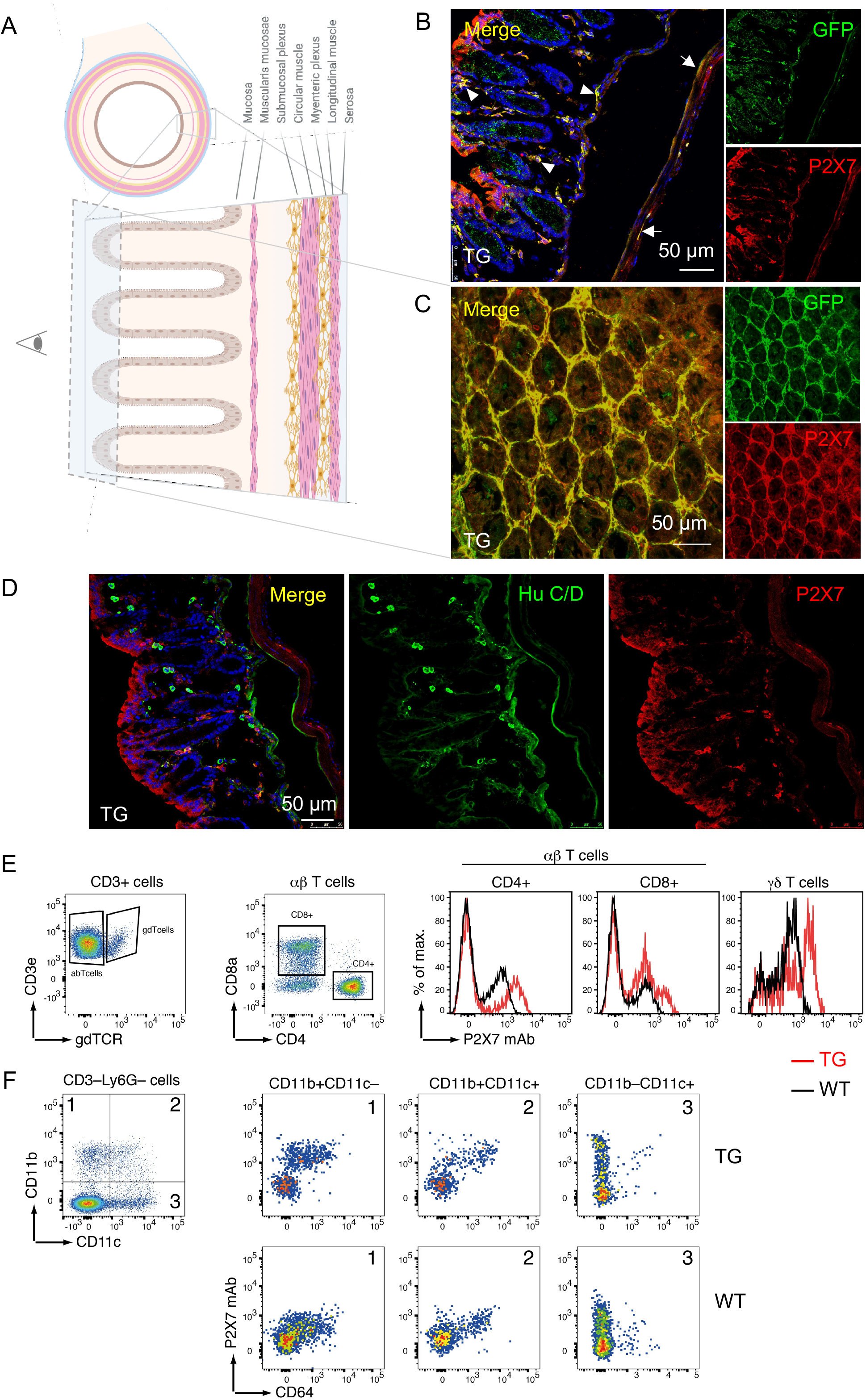
P2X7 localization in different layers of the mouse colon. (A) Schematic drawing (created with BioRender.com) of the histological structure of the colon explaining the cell layers and perspectives of the images shown in B-D. (B) Co-staining of colonic cross section and a planar mucosa preparation from P2X7-EGFP mice with anti-P2X7 and anti-GFP antibodies. Arrow heads and arrows in (B) indicate examples of single cells and layers with clear co-localization. (D) Co-staining of colonic cross section with anti-P2X7 antibodies and an antibody against the neuronal marker HuCD. All scale bars: 50 μm, cell nuclei were counterstained with DAPI (blue). (E) Flow cytometric analyses of colonic immune cells from wt (black) and transgenic (red) mice. Within the CD3^+^ T cell population, CD4^+^ and CD8^+^ αβ T cells and γδ T cells were analyzed for cell surface P2X7 expression using the mAb RH23A44. The right-shifted histogram indicates higher P2X7 expression in transgenic mice. (F) Comparison of P2X7 expression in innate immune cells. Among CD3-Ly6G-innate immune cells, CD11b^+^CD11c^−^, CD11b^+^CD11c^+^, and CD11b^−^CD11c^+^ cells were analyzed for cell surface P2X7 expression using mAb RH23A44. Cells were co-stained with anti-CD64.

To compare P2X7 expression in immune cells of the colon from P2X7-EGFP transgenic and WT mice, we performed flow cytometry and stained immune cells isolated from colon with an anti-P2X7 monoclonal antibody. Among T cells, CD4^+^, CD8^+^, and γδ T cells from P2X7-EGFP transgenic and WT mice exhibited a similar P2X7 expression pattern, though cells from P2X7-EGFP transgenic mice showed in general higher P2X7 level due to transgene overexpression (Fig. 1E). Similarly, colon-derived innate immune cells, differentiated by the expression of CD11b and CD11c, showed a comparable P2X7 expression pattern in P2X7-EGFP transgenic and WT mice. Here, P2X7 expression was prominent on CD11b+CD11c– and CD11b+CD11c+ cells co-expressing CD64+ (Fig. 1F).

### P2X7 expression in macrophages of the myenteric plexus

To further investigate P2X7 expression in the muscular layer and identify the respective P2X7-expressing cell types, we next stained whole-mount preparations of the longitudinal muscle and myenteric plexus (LMMP) from P2X7-EGFP, wt, and P2rx7^−/−^ mice with the P2X7-specific nanobody. This revealed specific staining of enteric ganglia that appeared as mesh-like structures and large isolated extraganglionar cells with ramified morphology (Fig. 2A, compare also Figs. 3A and 5A). Staining of the latter was particularly strong in the P2X7-EGFP mice, which was confirmed by co-staining with an antibody against GFP (Fig 2B). In contrast, the EGFP signal in the mesh-like structures was less consistent and had a patchy appearance, suggesting that EGFP was only present in some clusters of the cells that are forming this structure (compare two representative images in Fig. 2B). Co-staining with antibodies against F4/80 (EGF-like module-containing mucin-like hormone receptor-like 1), Iba1, and CD68 confirmed that the extraganglionar cells represent macrophages, in agreement with the known expression of P2X7 receptors in these cell types (Fig. 2C). Two different types of macrophages could be differentiated: macrophages with longitudinal orientation and bipolar morphology likely represent muscular macrophages whereas those with ramified morphology were localized in the vicinity of ganglions. Interestingly, only the ramified macrophages were positive for Iba1 (see also Suppl. Fig. 1). Since macrophage numbers appeared to be increased in P2X7-EGFP mice in comparison to wt mice (Fig. 2 D, compare also Fig. 3A), we performed a quantitative analysis by counting the F4/80-positive macrophages in randomly chosen images of both genotypes. This revealed about two times more macrophages in the myenteric plexus of P2X7-overexpressing mice (97.03 ± 12.91 and 202.7 ± 32.03 macrophages per image (0.125 mm^2^, n=5 images/mouse) in WT (N=3) and transgenic mice (N=4), respectively). Finally, co-stainings with antibodies against Anoctamin-1 (ANO-1) and CD117 excluded the presence of P2X7 receptors in myenteric interstitial cells of Cajal that function as pacemaker cells (Suppl. Fig. 2). The good correlation between stainings of ANO-1 and CD117, which represents also a mast cell marker, is in agreement with a low number of mast cells in the myenteric plexus.

**Figure 2:**
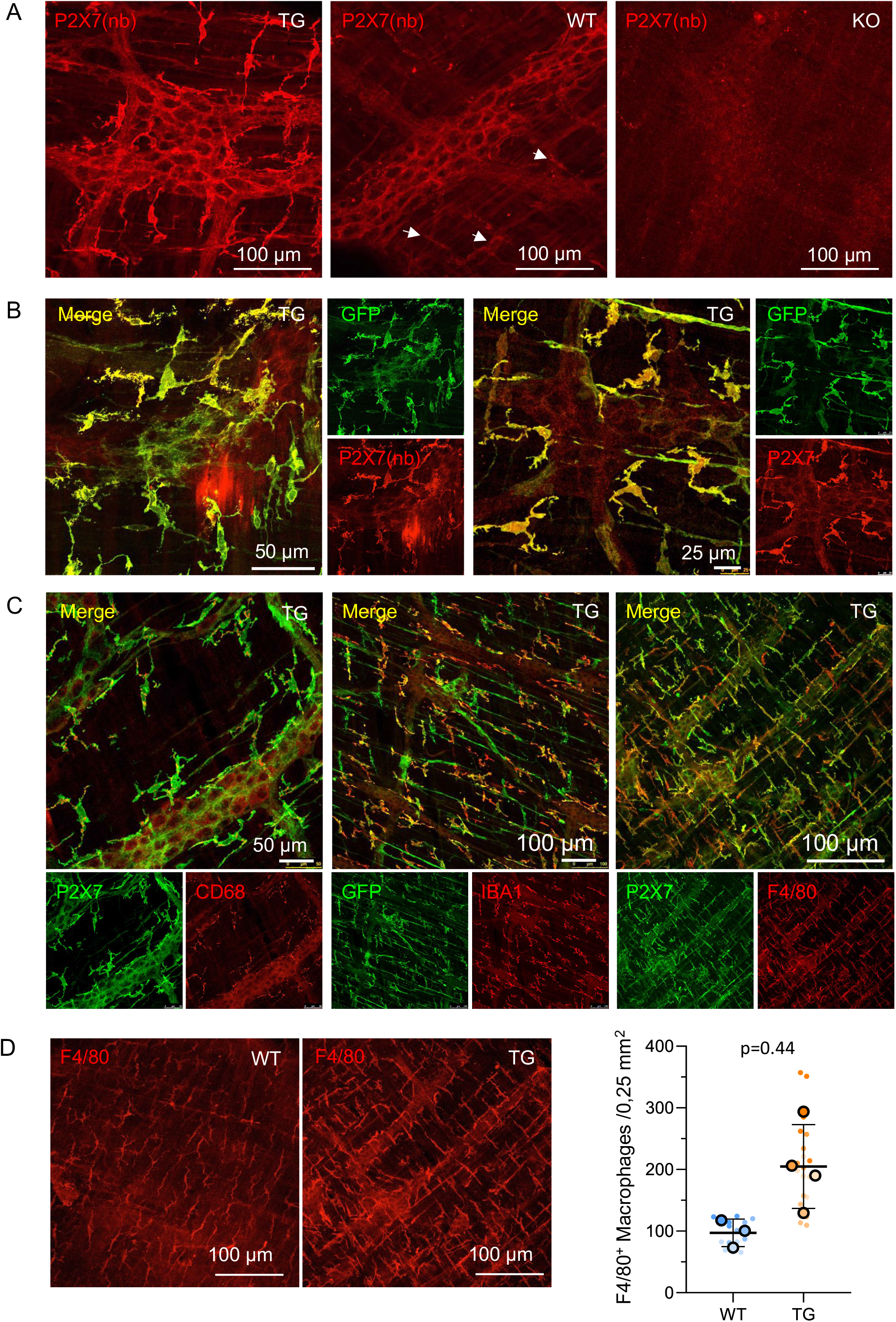
P2X7 staining in whole mount. LMMP preparations. (A) Specificity of the P2X7-specific nanobody. Preparations from P2X7-EGFP, wt, and *P2rx7^−/−^* mice were stained as indicated and confocal images were taken with identical settings. Arrows indicate extraganglionar P2X7 positive cells. (B) Two representative confocal images of preparations from P2X7-EGFP transgenic mice co-stained with anti-GFP and anti-P2X7 antibody or nanobody, as indicated. (C) Tissues from P2X7-EGFP mice were co-stained with anti-P2X7 or anti-GFP antibodies and antibodies against different macrophage marker proteins. (D) Representative images of tissue from P2X7-EGFP and wt mice stained with an antibody against the macrophage marker F4/80 and statistical analysis of macrophage numbers in both genotypes. n=5 images of 0.125 mm^2^ from the distal colon were analyzed per mouse. Data are represented as mean ± SEM from N = 3 - 4 mice per group (large symbols). Numbers for individual mice are represented in small symbols with different color intensities for each mouse. 97.03 ± 12.91 and 202.7 ± 32.03 macrophages per image were counted in WT and transgenic mice, respectively. Statistical significance was tested by Welch’s t-test (p=0,0438). Normal distribution (Shapiro-Wilk, Kolmogorov-Smirnov, QQ plot) was confirmed for each group.

**Figure 3:**
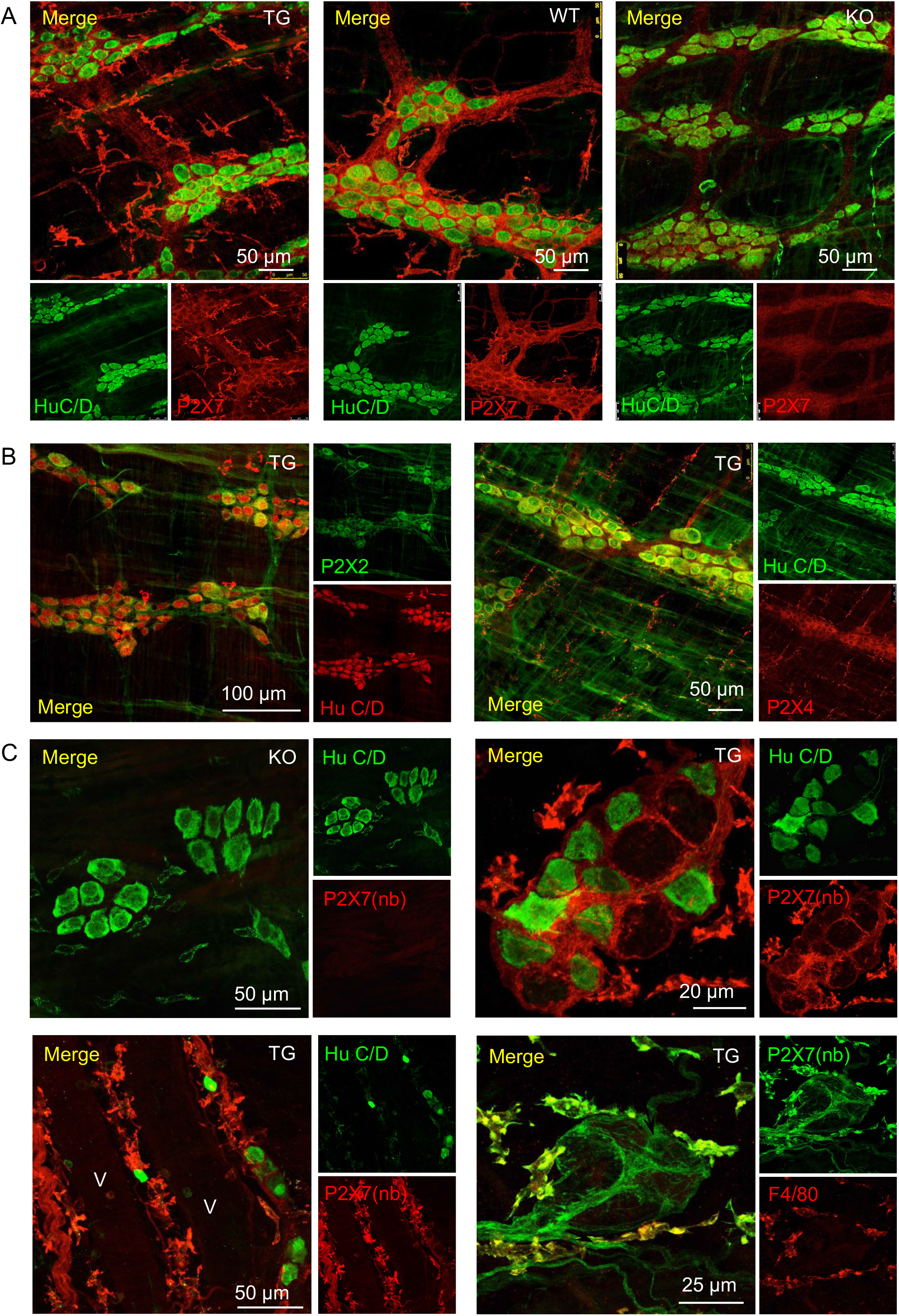
Analysis of neuronal P2X7 localization in LMMP and submucosal plexus preparations. **(A)** Specificity of staining with the P2X7-specific antibody in combination with antibodies against the neuronal marker HuC/D. LMMP preparations from P2X7-EGFP, wt, and *P2rx7^−/−^* mice are shown. (B) Co-staining of HuC/D and P2X2 receptors or P2X4 receptors (both used as positive controls for neuronal P2X expression) in LMMP preparations from P2X7-EGFP mice. (C) Co-staining of P2X7 (using the P2X7 nanobody) and HuC/D in submucosal plexus preparations of *P2rx7^−/−^* and P2X7-EGFP mice, as indicated (upper panel). The images from P2X7-EGFP mice demonstrate staining of ganglia (right panel) and blood vessels (lower left image). V indicates the lumen of the blood vessel. The lower right image confirms P2X7 co-staining with the macrophage marker F4/80.

### No evicence for P2X7 expression in neurons of the myenteric and submucosal plexuses

We next performed co-staining of HuC/D and P2X7 in the myenteric plexus (Fig. 3A and Suppl. Fig. 3) and the submucosal plexus (Fig. 3C) to examine a possible neuronal localisation of P2X7. These stainings revealed a clearly different localization of HuC/D and P2X7 in both wt and P2X7-EGFP mice. Similar results were obtained if co-stainings were performed using the anti-P2X7 nanobody or the anti-GFP antibody for P2X7 staining (Suppl. Fig. 3). As positive control for a neuronal expression pattern, we used antibodies against P2X2 and P2X4 receptors, which were previously shown to be expressed in enteric and other neurons (Galligan and North, 2004; Linan-Rico et al., 2015; Nieto-Pescador et al., 2013; Ren et al., 2003). Accordingly, these stainings revealed good co-localization of HuC/D with both receptors in neuronal cell bodies (Fig. 3B). While a uniform staining in all neurons was seen for the P2X4R, P2X2R staining appeared more variable between cells. In contrast to P2X2Rs, a punctate P2X4R staining was also detected in extraganglionic cells. Here P2X4 co-localized with CD68, in agreement with its known expression in lysosomes of macrophages (Suppl. Fig. 4), (Duveau et al., 2020).

HuC/D is localized in the cell body but under pathophysiological conditions, has been reported to show a prevalent nuclear localization (Desmet et al., 2014). Thus, to exclude P2X7 expression in specific neuronal substructures that are not stained with HuC/D, we also performed co-localization studies of P2X7 with β3-tubulin, which stains neuronal processes, the neuronal marker proteins PGP9.5 (protein gene product 9.5), calbindin and calretinin (Fig. 4A) as well as with the synaptic marker proteins synaptophysin, and vesicular glutamate transporter subtype 2 (vGlut2) (Fig. 4B). Again, none of these stainings revealed a co-localization with P2X7, thus arguing against a physiological expression of P2X7 protein in enteric neurons.

**Figure. 4:**
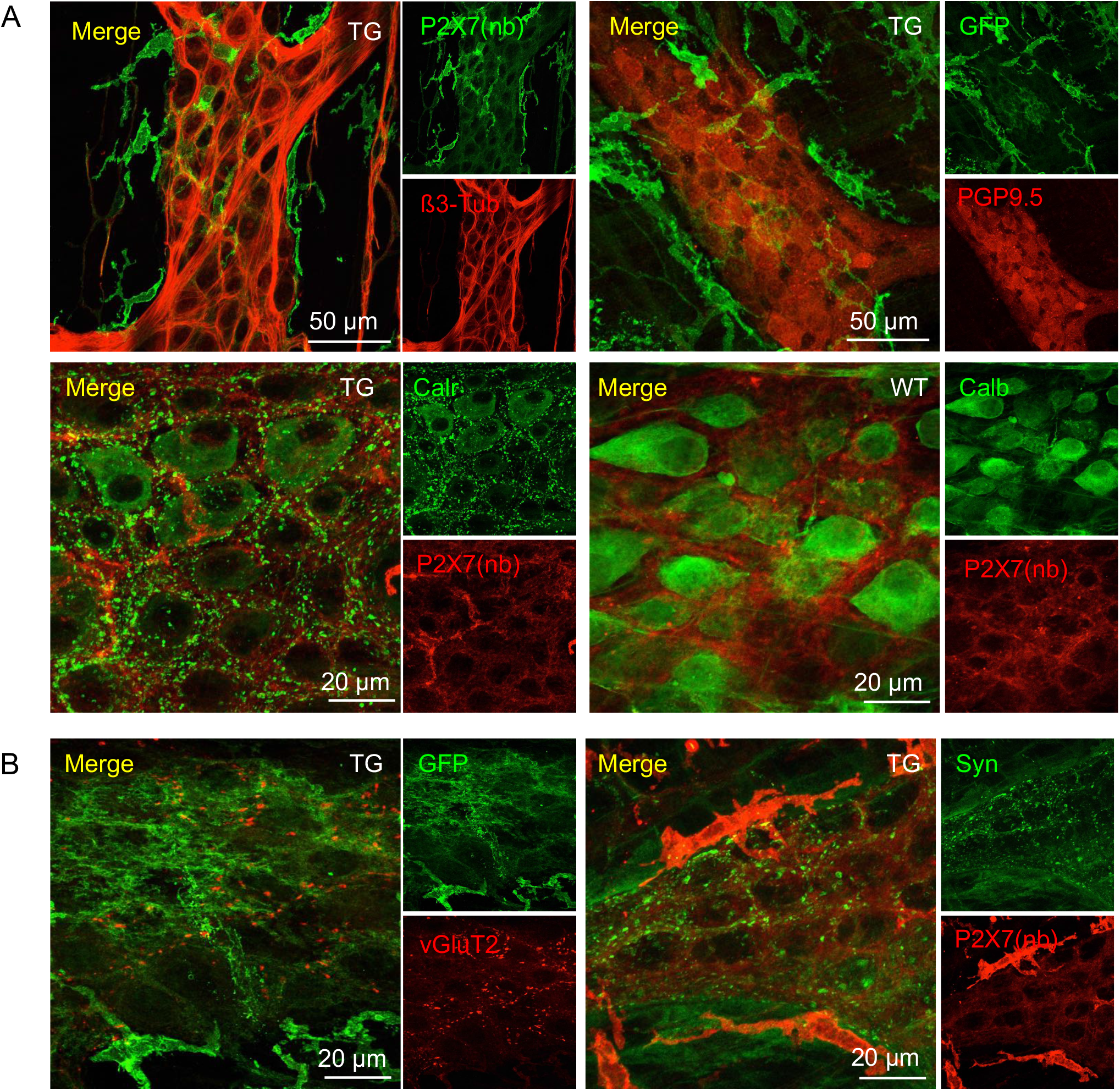
Analysis of neuronal and synaptic P2X7 localization in whole mount LMMP preparations. Tissues from P2X7-EGFP and wt mice were co-stained with P2X7-specific nanobody or anti-GFP antibodies as indicated and antibodies against different neuronal (A) or synaptic (B) marker proteins. β3-Tub, β3-tubulin; PGP9.5, protein gene product 9.5; Calr, Calretinin; Calb, Calbindin; vGluT2, vesicular glutamate transporter 2; Syn, synaptophysin.

Interestingly, the strong staining of the P2X7-EGFP-expressing macrophages enabled the visualization of close interactions of macrophages with the HuC/D-positive neurons (Fig. 4A), as previously described (Gabanyi et al., 2016; Muller et al., 2014; Phillips and Powley, 2012). In addition, a close association of macrophages with blood vessels in the submucosal plexus could be observed (Fig. 3C, right panel).

### Identification of P2X7 expression in enteric glia cells

Based on the above data, we concluded that the intraganglionic P2X7-expressing cells that are forming the mesh-like structure in which the neurons are embedded represent most likely enteric glia cells. Enteric glia cells are generally differentiated by their localization and morphology whereas expression of specific marker proteins appears highly dynamic (Boesmans et al., 2015; Gulbransen and Sharkey, 2012). Using antibodies against the widely used glia marker GFAP (glial fibrillary acidic protein) and S100β (S100 calcium-binding protein B), we detected a clear and specific co-localization of P2X-EGFP and S100β in wt and transgenic mice (Fig. 5). However, while a uniform intraganglionic and interganglionic staing of S100β is seen, the anti-GFP antibody shows a patchy pattern, in agreement with the incomplete EGFP P2X7 co-staining in these structures (compare also Fig. 2B above).

**Figure 5:**
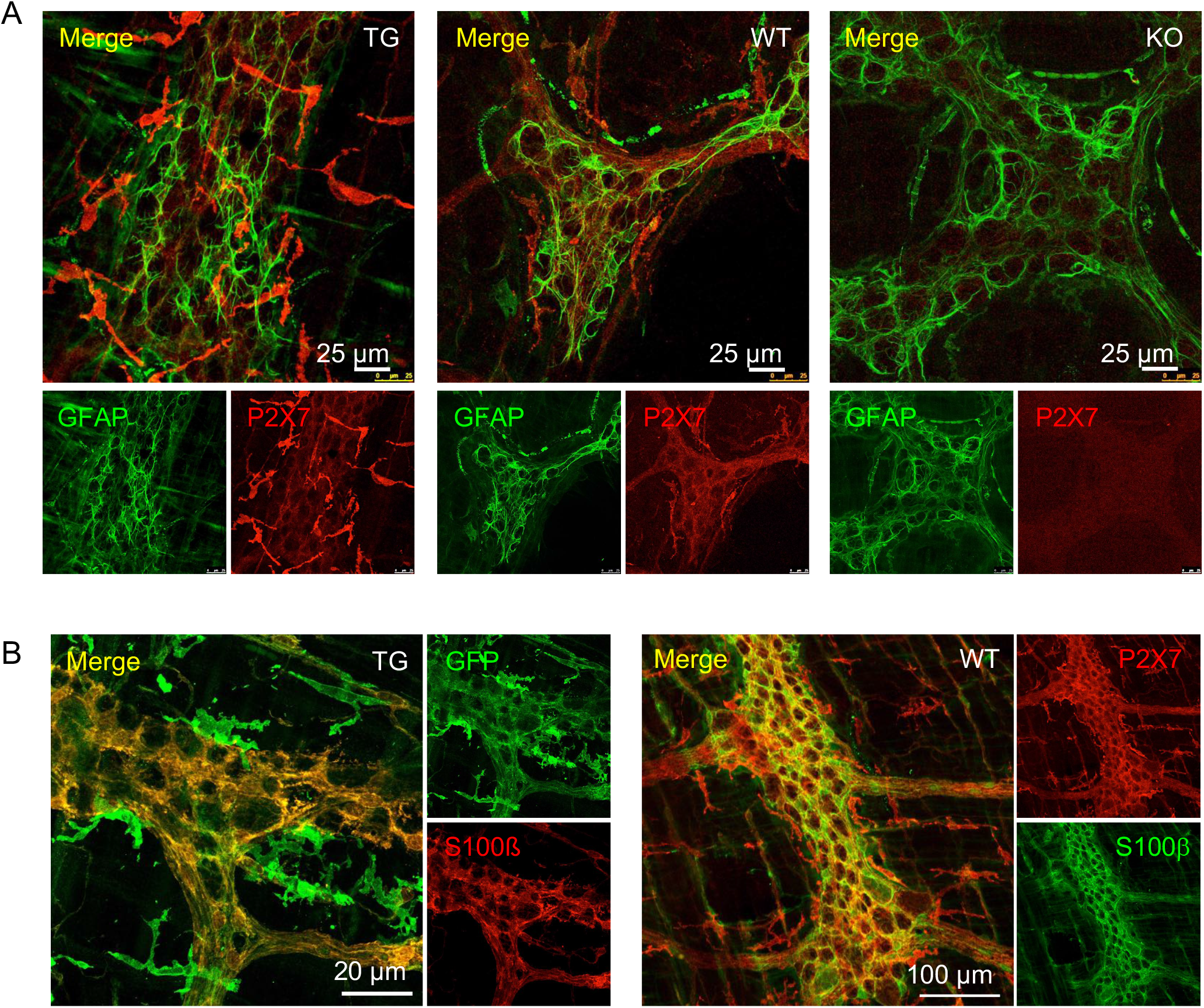
Analysis of P2X7 localization in enteric glia of the myenteric plexus. **(A)** Specificity of staining with the P2X7-specific antibody in combination with antibodies against the glial marker GFAP. LMMP preparations from P2X7-EGFP, wt, and *P2rx7^−/−^* mice are shown. (C) Co-staining of P2X7-EGFP (using the anti-GFP antibody) and the glial marker S100β in tissue from P2X7-EGFP mice. Two representative images are shown to demonstrate the patchy distribution of P2X7-EGFP expression in enteric glia.

### No induction of neuronal P2X7 expression in the DSS colitis model

The above data show a dominant expression of P2X7 in macrophages and glia cells but not in enteric neurons. This is in contrast to previous studies, where neuronal P2X7 receptors have been implicated in colitis-associated enteric neuron death (Gulbransen *et al*., 2012). We therefore wondered, whether neuronal P2X7 expression could be induced by Bz-ATP application in an *ex vivo* model in which we incubated *myenteric plexus* preparations from wt, P2X7-EGFP and *P2rx7^−/−^* mice for 60 min with 300 μM Bz-ATP. In agreement with previous observations (Gulbransen *et al*., 2012), Bz-ATP-treatment resulted in a reduction of neuron numbers. This appeared to be increased in transgenic mice, although the difference was not significant (Fig. 6A). Most importantly, no upregulation of P2X7 expression in neurons was observed, neither in wt, nor in transgenic mice. To examine a possible neuronal P2X7 upregulation under more physiological conditions and investigate if P2X7 overexpression is associated with a higher disease susceptibility, we next subjected wt and transgenic mice to the DSS colitis model. Figures 6B and C show only a slightly increased weight loss of DSS-treated P2X7-EGFP mice compared to DSS-treated wt mice at day 10, but no effects on the fecal score during the entire experiment and colon length at day of sacrifice. Microscopic analysis of HuC/D-stained tissue from DSS-treated mice shows massive tissue damage but no neuronal P2X7 localization (Fig. 6D).

**Figure 6:**
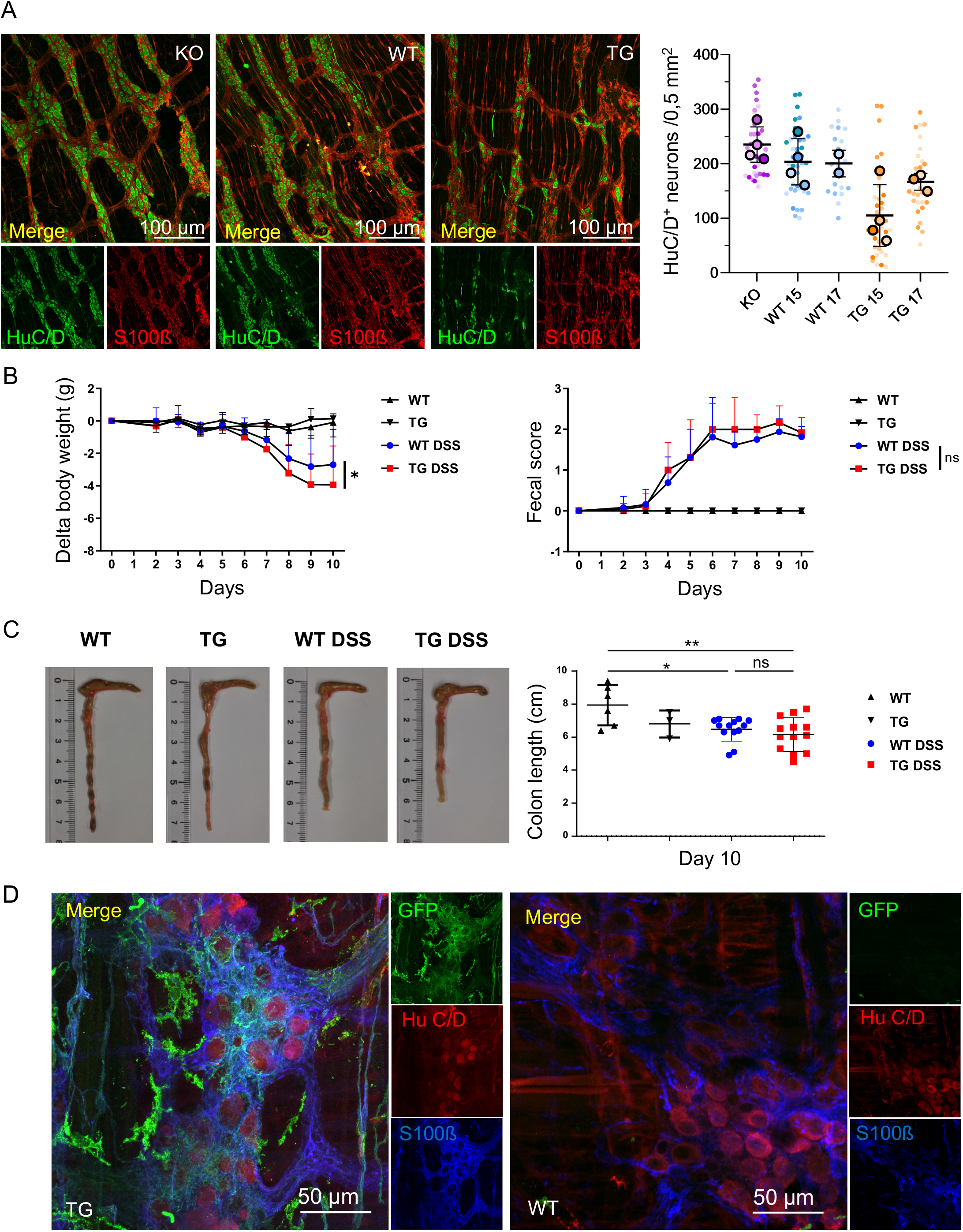
Comparison of wt and P2X7-EGFP overexpressing mice in an *ex vivo* model and in the *in vivo* DSS colitis model. (A) Representative images of LMMP preparations after treatment with 300 μM Bz-ATP (line 15) and statistical analysis of neuron numbers per 0.501 mm^2^. Data from knockout mice, and transgenic P2X7-EGFP lines 15 and 17, and their respective wildtype litter mates were analyzed by Brown-Forsythe and Welch ANOVA with a post-hoc Tamhane T2 test to correct for inhomogeneity of variance. Data are represented as mean ± SEM from N = 2 - 4 mice per group (large symbols). Numbers from n = 9 - 10 images per mouse are represented in small symbols with different color intensities for each mouse. (B) Delta of weight variation (left) and fecal score (right) over time in P2X7-EGFP (TG, line 17) mice and their wild-type control littermates (WT) treated or not with DSS 1%. (C) Pictures (left) and statistical analysis of colon length (right) at day 10 in P2X7-EGFP (TG) mice and their wild-type control littermates (WT) treated or not with DSS 1%. ns: not significant; *: P < 0.05; **: P < 0.01. Fecal score: 0, normal solid faeces; 1, soft faeces; 2, liquid faeces; 3, visible blood traces on faeces. Data are represented as mean ± SD. All mice were in C57/BL6 background. Female mice of 16 weeks of age were used in (B) and (C). (D) Triple staining of tissue from DSS-treated wt and P2X7-EGFP transgenic mice with antibodies against GFP, the neuronal marker HuC/D and the glial marker S100β.

## Discussion

Despite its importance as a drug target, the localization, regulation, and functions of P2X7 receptors in different cell types remains controversial. This is partly due to the complex pharmacology of purinergic receptors and the unclear specificity of the available antibodies in different tissues. In the study presented above, we analysed the cell type-specific P2X7 localization in the colon with a focus on the myenteric plexus.

We demonstrate that myenteric macrophages and enteric glia represent the dominant P2X7-expressing cell types in the myenteric plexus under both physiological and pathological conditions and visualize the close interaction of the macrophages with enteric neurons. Based on our data we propose that P2X7 receptors in macrophages and/or glia cells indirectly contribute to neuronal damage in colitis pathology.

### P2X7 in enteric neurons and glia

The enteric nervous system and enteric glia cells have emerged as important regulators of gut barrier function and homeostasis in gastrointestinal diseases including IBD (Sharkey et al., 2018). P2X7 receptors have been described in enteric neurons (da Silva *et al*., 2017; Palombit *et al*., 2019)(Antonioli *et al*., 2014; Brown *et al*., 2016; Gulbransen *et al*., 2012)(Evangelinellis *et al*., 2021) and a role of neuronal P2X7 receptors in neuronal cell death and colitis-associated motility disorders was proposed. However, using antibodies against a variety of neuronal marker proteins in distinct neuronal substructures in combination with P2X7-specific anti- and nanobodies and *P2rx7^−/−^* mouse controls as well as a transgenic P2X7-EGFP reporter mouse model, we could not detect neuronal P2X7 receptors. In contrast, a clear staining was found with P2X2 and P2X4-specific antibodies that were used as positive controls. Based on these data we conclude that P2X7 protein is either absent in neurons or below the detection limit and that P2X7 receptors do not play a physiological role in enteric neurons.

### Presence and function in glia cells

Enteric glia are closely associated with enteric neurons. They modulate motility, secretion, function of the epithelial barrier, neuroinflammation, and have also been involved in nociceptor sensitization and immune cell modulation (Grubisic et al., 2020). Using both an antibody and a nanobody against P2X7, we could detect identical patterns of P2X7 expression that were absent in *P2rx7^−/−^* mice and correlated well with the expression of the enteric glia cell marker S100β. Since the S100β antibody and both the P2X7-specific antibody and nanobody carry rabbit Fc domains, no co-staining could be performed for these proteins. Using the P2X7-EGFP reporter mouse, however, the co-localization of P2X7-EGFP and S100β could be confirmed although P2X7-EGFP expression appeared more irregular than that of the endogenous P2X7. The reason for this is unclear but might reflect the highly dynamic protein expression and subtype differentiation that has been reported for enteric glia (Boesmans *et al*., 2015; Gulbransen and Sharkey, 2012). In support of our finding, glial responses to ATP and expression of P2X7 in enteric glia was previously reported in rats (da Silva *et al*., 2017; Mendes et al., 2019; Van Crombruggen et al., 2007; Vanderwinden et al., 2003) and a glial cell line (Bhave et al., 2017). However, purinergic neuron to glia signaling and ATP-mediated Ca^2+^ signals in enteric glia cells were generally believed to be mediated by P2Y1 receptors (Boesmans *et al*., 2015; Gulbransen and Sharkey, 2012) (Boesmans et al., 2013; Gomes et al., 2009). Thus, while the importance of glia in neuro-immune interaction in colitis associated visceral pain has recently been shown (Grubisic *et al*., 2020), the role of glial P2X7 receptors remains completely unclear. Interestingly, enteric glia in the myenteric plexus share marker proteins and functional properties with satellite glia in the DRG and astrocytes in the CNS and common roles in inflammatory processes and abdominal pain generation were described (for a review see (Morales-Soto and Gulbransen, 2019)). Both satellite glia and Bergmann glia, a specific type of S100β-positive astrocytes, express also P2X7, suggesting common roles of P2X7 receptors in these cells. Further studies are needed to clarify the functions of glial P2X7 receptors and their involvement in colitis pathology and visceral pain.

### Presence and function in macrophages

Based on their localization and specific functions, intestinal resident macrophages are classified into two major groups: the highly phagocytic lamina propria macrophages that represent the first line of defence against pathogens and act primarily as innate immune effector cells and the muscularis macrophages, which represent the dominant immune cell population in the muscular layer and closely and bidirectionally interact with the enteric nervous system and also with smooth muscle cells (Viola and Boeckxstaens, 2020). Although of different developmental origin, macrophages share many marker proteins with microglia, the resident immune cells of the CNS (Ginhoux and Prinz, 2015). Muscularis macrophages resemble microglia also morphologically and functionally (Verheijden et al., 2015). P2X7 receptors are highly expressed in microglia and macrophages and staining of overexpressed P2X7-EGFP with P2X7-specific nanobodies and antibodies or antibodies against GFP resulted in a clear and detailed visualization of macrophages and revealed their close interaction with neuronal cell bodies and processes, similar to microglia in the CNS. The P2X7-EGFP mice might therefore provide a good model to further study these interactions. As the maturation and release of cytokines, such as IL-1β, is the best investigated function of P2X7 receptors in macrophages, this represents the most obvious mechanism how P2X7 could contribute to inflammation but also nociceptor sensitization (Grubisic *et al*., 2020). P2X7-mediated production of reactive oxygen species (Kopp et al., 2019) could further contribute to ATP-induced neuronal damage or death as observed in (Gulbransen *et al*., 2012). Since intestinal macrophages, similar to microglia in the CNS, are considered gatekeepers of tissue homeostasis, they are also discussed as therapeutic targets in IBD (Na et al., 2019). The physiological mechanism or relevance of the increased macrophage number in P2X7-EGFP mice remains to be determined but its reminiscent of a trend towards higher microglia numbers that was found in a preliminary analysis of the retina of P2X7-EGFP mice (Kaczmarek-Hajek *et al*., 2018). An intriguing possibility would be that P2X7 overexpression has an effect on macrophage migration and/or proliferation.

### Involvement of P2X7 in other cell types

P2X7 receptors in other cell types besides macrophages and glia have also been involved in colitis pathology. Thus, P2X7 receptors are highly expressed in mast cells and P2X7-mediated mast cell activation was shown to initiate and exacerbate colitis via cytokine induction and neutrophil infiltration (Kurashima *et al*., 2012). In support of this finding, inhibition of ATP-induced mast cell activation ameliorated colitis (Matsukawa *et al*., 2016; Ohbori *et al*., 2017). P2X7 receptors have also been involved in T cell activation and its expression is dynamically regulated during T cell differentiation. T cell subsets, including follicular helper T cells, regulatory T cells and Th17 T cells, are sensitive to P2X7-mediated cell death (Proietti *et al*., 2014; Rissiek *et al*., 2015) and it was reported that P2X7-mediated death of regulatory T cells decreases immune tolerance and thereby promotes inflammation (Figliuolo *et al*., 2017; Schenk *et al*., 2011). In epithelial cells, activation of P2X7 receptors was shown to induce IL-1β release (Cesaro *et al*., 2010) and thereby, suggested to promote inflammation. Furthermore, it was shown to inhibit epithelial cell proliferation and P2X7 receptor inhibition or deletion promoted re-epitheliation and recovery of inflammatory lesions but also enhanced tumor susceptibility (Hofman *et al*., 2015). Based on a study on human colonic mucosal strips, it was suggested that epithelial P2X7 receptors represent therapeutic targets for IBD (Diezmos *et al*., 2018).

In conclusion, P2X7 receptors in numerous cell types appear to contribute to colitis pathology as well as associated motility disorders and visceral sensitivity. Here we demonstrate that in the intestinal nervous system, P2X7 receptors are dominantly expressed in macrophages and S100β-positive glia. This is resembling the situation in the CNS, where P2X7 receptors are highly expressed in microglia and S100β-positive astrocytes. We propose that P2X7 fulfills similar functions in both systems and that ATP-induced neuronal damage in the enteric system is an indirect effect of macrophage or glial P2X7 activation. Since numerous studies support beneficial effects of P2X7 blockade in central and peripheral inflammatory diseases it represents a promising drug target. and this study provides a basis to further elucidate its cell type-specific functions and develop novel therapies.

## Author contributions

Conceptualization: AN, TJ, JZ, SZ

Formal analysis: TJ, JZ, TRJ, BR, SZ, MS, FG, AN, TM

Investigation: TJ, JZ, TRJ, BR

Resources: FKN

Writing - Original Draft: AN

Writing-Review and Editing: TJ, JZ, TRJ, SZ, MS, FG, AN, BR, TM, FKN

Project administration: AN

Funding acquisition: AN, MS, FG, TM, FKN

## Acknowledgements

This work was supported by funding from the Förderprogramm für Forschung und Lehre (FöFoLE) of the LMU (No. 745/2014), and the Deutsche Forschungsgemeinschaft (DFG, German Research Foundation, Project-ID: 335447717 - SFB 1328 and TRR-152, P14) and the Swiss National Science Foundation (Project number IZCNZ0-174704).

We thank Barbara Kuch for technical assistance, Béla Zimmer for help with microscopy, and Yves Haufe for statistical advice.

## Material and Methods

### Animals

BAC transgenic P2X7-EGFP mice (Tg(RP24-114E20P2X7451P-StrepHis-EGFP)Ani15 and 17) were recently described (Kaczmarek-Hajek *et al*., 2018) and in FVB/N or C57BL6 background. Identical P2X7 expression patterns were observed in both lines and strains. *P2rx7^−/−^* mice (P2rx7^tmid(EUCOMM)Wtsi^) were bred as described (Kaczmarek-Hajek *et al*., 2018) and in C57BL/6 background. 12-18 weeks old animals of both sexes were used. Mice were housed in standard conditions (22°C, 12 h light– dark cycle, water/food ad libitum). All animal experiments were performed in accordance with the principles of the European Communities Council Directive (2010/63/EU). Procedures were reviewed and approved by the State of Upper Bavaria (55.2-1-54-2532-59-2016). All efforts were made to minimize suffering and number of animals.

### Flow cytometric analysis

For purification of leukocytes from the large intestine, the colon was separated from the small intestine, opened by longitudinal section and feces was washed out thoroughly in Hanks Balanced Salt Solution (HBSS). The cleaned colon was cut into 1 cm pieces and further washed in 20 ml HBSS at 37°C for 30 min. Colon pieces where then digested for 30 min at 37°C in a digestion solution containing collagenase (1 mg/ml) and DNase (0.1 mg/ml). The generated cell suspension was filtered through a 100 μm cell strainer and centrifuged for 5 min at 300 g. Leukocytes were separated from debris by resuspending the pellet in 5 ml 33% Percoll solution (GE Healthcare). Erythrocytes were lyzed using an ACK lysis buffer (155 mM NH_4_Cl, 10 mM KHCO_3_, 0.1 mM EDTA, pH 7.2). Isolated cells were stained with an antibody mix in FACS buffer (containing 1 mM EDTA (Sigma) and 0.1 % bovine serum albumin (Sigma)) for 30 min at 4 °C (antibodies see key resource table) to identify major immune cell populations, including T cells and innate immune cells. Cells were washed once with FACS buffer and were subsequently analyzed on a Becton Dickinson FACSymphony A3 cell analyzer.

### Whole mount myenteric plexus and submucosal plexus preparation

For whole mount myenteric plexus and submucosal plexus preparation, mice were sacrificed by cervical dislocation and 1 cm segments were taken 1 cm aboral from the proximal colon and transferred to ice-cold Krebs solution (containing in mM: 117 NaCl, 4.7 KCl, 1.2 MgCl_2_, 1.2 NaH_2_PO_4_, 25 NaHCO_3_, 2.5 CaCl_2_, 11 glucose, aerated with carbogen to pH 7.4) in Sylgard (Dow Corning)-filled dissecting dishes. After flushing with Krebs buffer, segments were opened along the mesenteric border, pinned out, and fixed for 4 hr at 4°C (4% PFA and 0.2% picric acid in 0.1 M phosphate buffer (pH 7.4). Tissue was rinsed (3 × 10 min) with phosphate buffer and dissected in PBS (pH 7.4). Using forceps and a stereomicroscope, the mucosa and submucosa, were carefully removed and the submucosal plexus was then transferred to a microscopic slide. For myenteric plexus preparations, the circular musculature was also removed. Preparations were blocked (0.5%, Triton X-100, 0.1% NaN_3_, 4% goat serum (Sigma-Aldrich) in PBS, 1 hr, RT), incubated with primary antibodies in the above solution (12 hr at RT), rinsed 3 × 10 min with PBS, and incubated with secondary antibodies (2 hr at RT). After washing (3x in PBS) preparations were mounted in PermaFluor (Thermo Fisher Scientific) on slides and images were acquired with a Zeiss 880 Airyscan confocal microscope and processed with ImageJ.

### Preparation and staining of colon cryo-sections

Animals were anesthetized with isoflurane (IsoFlo®) and euthanized by cervical dislocation. The distal colon was prepared, flushed 3 times with PBS, embedded in Tissue Tek® and after freezing at −80°C cut in 10 μm cross-sections using a Cryostat Microm HM560 (Leica Biosystems).

For immunofluorescence staining, colon sections were dried for 1 h at RT, fixed for 20 min (2% PFA in 150 mM sodium phosphate buffer pH 7,4), and washed 2 x 10 min with PBS and 1 x 10 min with water. After drying at RT, slices were blocked for 1 h (10% goat serum, 0,5% Triton X-100, in PBS, pH 7,4) and then incubated overnight at 4°C with 100 μl of the primary antibody dilution in blocking solution using a humid chamber.

After washing (3x 10 min in PBS), 100 μl of the secondary antibody dilution in blocking buffer were applied for 1 h at RT and slices were then washed (2 x 10 min with PBS, 1 x 10 min with water). Sections were subsequently treated for 1 min with 0.1% DAPI, washed 3 x 5 min with PBS, and embedded in PermaFluor™ Mounting Medium. Preparations were analyzed using a Zeiss 880 Airyscan confocal microscope..

### Counting analysis

For counting analysis, 1cm^2^ whole-mount LMMP preparations taken 1 cm aboral from the beginning of distal colon were used. 5 and 10 z-stack images (0.5 mm^2^) were randomly selected for counting of macrophages or neurons, respectively. Only F4/80-positive macrophages or HuCD-positive neurons that were completely within the image were counted using the image J point tool.

### Bz-ATP treatment

For *ex vivo* Bz-ATP treatment of LMMP preparations a modified protocol of (Gulbransen *et al*., 2012) was used. The distal colon was prepared and fixed with minutien pins in a Sylgard dish, and opened along the mesenteric border as described above. Mucosa, submucosa, and plexus submucosus were prepared off the muscle layer. The circular muscle layer was maintained to avoid shearing stress of the MP. The tissue was then incubated for 1 h in a cell culture incubator with 20ml of 300 μM BzATP in Krebs buffer (containing in mM: 117 NaCl, 4.7 KCl, 1.2 MgCl_2_, 1.2 NaH_2_PO_4_, 25 NaHCO_3_, 2.5 CaCl_2_, 11 glucose, aerated with carbogen to pH 7.4). The tissue was then washed 3 x 10 min with Krebs buffer and incubated for another 2 h with 20 ml Krebs buffer before fixation for 4 h in 4% PFA (4% PFA and 0.2% picric acid in 0.1 M phosphate buffer (pH 7.4) at 4°C. Subsequently, the circular muscle layer was removed and immunofluorescence staining was performed as described above.

### DSS colitis model

BAC transgenic P2X7-EGFP mice (Tg(RP24-114E20P2X7451P-StrepHis-EGFP)Ani17) (Kaczmarek-Hajek *et al*., 2018) and their wild-type (wt) control littermates were maintained and treated in accordance with the Swiss Federal Veterinary Office guidelines at IRB animal facility. Experiments were approved by ’’Dipartimento della Sanitá e Socialitá’’ with authorization numbers TI-25/2018 and TI-24/2019. For experimental procedures 16 weeks old female mice were used. Mice were housed in standard conditions (20-22°C, 12 h light–dark cycle, water/food ad libitum) and monitored and weighted daily. The following values were used to establish the daily fecal score: 0, normal solid faeces; 1, soft faeces; 2, liquid faeces; 3, visible blood traces on faeces. All efforts were made to minimize suffering and limit the number of animals used in experiments.

Dextran sodium sulphate (DSS, TdB Labs) was added at a concentration of 1% in drinking water for 6 days, then DSS 1% was replaced with normal drinking water for other 4 days. After 10 days from the beginning of the experiment, mice were sacrificed by inhalation of CO2 for retrieval of organs.

### Statistical analysis

Graph Pad Prism software was used for statistical analysis and data were presented as means ± standard derivation (SD). Student’s t- test, one way or two way ANOVA (Turkey’s multiple comparisons test) was used to determine statistical differences between groups. Significance was accepted at * p < 0.05, ** p < 0.01 *** p < 0.001.

Key Resource Table: Antibodies used for immunohistochemistry and flow cytometry.

| **Antibody** | **Supplier** | **Cat.# and RRID** | **Dilution** |
| --- | --- | --- | --- |
| GFP<br>(rb pAb) | Abcam | ab6556<br>AB_305564 | 1:2000 |
| GFP<br>(chk pAb) | Thermo Fisher | CA10262<br>AB_2534023 | 1:200 |
| P2X7 C-term<br>(rb pAb) | Synaptic<br>Systems | 177003<br>AB_887755 | 1:500 |
| P2X7 ECD<br>7E2-rbIgG | Nolte lab | Nanobody rbIgG fusion<br>construct | 0.1 µg/mL |
| P2X2<br>(rb pAb) | Alomone | APR-003<br>AB_2040054 | 1:200 |
| P2X4<br>(rb pAb) | Alomone | APR-002<br>AB_2040058 | 1:200 |
| Iba1<br>(rb pAb) | WAKO | 019-19741<br>AB_839504 | 1:500 |
| CD68<br>(rat FA-11) | Biorad | MCA1957<br>AB_322219 | 1:200 |
| F4/80<br>(rat BM8) | Thermo Fisher | 14-4801-81<br>AB_467557 | 1:100 |
| Hu C/D<br>(ms 16A11) | Thermo Fisher | A-21271<br>AB_221448 | 1:200 |
| β3-Tubulin<br>(gp pAb) | Synaptic<br>Systems | 302304<br>AB_10805138 | 1:200 |
| PGP9.5<br>(rb pAb) | Acurate Chemical<br>& Scientific Corp. | UCRA95101<br>AB_2313685 | 1:30.000 |
| Calretinin<br>(ms 37C9) | Synaptic<br>Systems | 214111<br>AB_2619904 | 1:200 |
| Calbindin D28k<br>(ms CB-955) | Sigma-Aldrich | C9848<br>AB_476894 | 1:200 |
| vGlut2<br>(rb pAb) | Synaptic<br>Systems | 135 403<br>AB_887883 | 1:500 |
| Synaptophysin<br>(ms 7.2) | Synaptic<br>Systems | 101011<br>AB_887824 | 1:500 |
| GFAP<br>(ms pAb) | Millipore | MAB360<br>AB_11212597 | 1:200 |
| S100β<br>(rb pAb) | Synaptic<br>Systems | 287003<br>AB_2620024 | 1:200 |
| Ano-1<br>(rb pAb) | Alomone | ACL-011<br>AB_10920576 | 1:50 |
| CD117<br>(rat ACK2) | eBioscience | 14-1172-82<br>AB_467436 | 1:1.000 |
| A594<br>gt anti-rb | Thermo Fisher | A11037<br>AB_2534095 | 1:400 |
| A594<br>gt anti-ms | Thermo Fisher | A11032<br>AB_2534091 | 1:400 |
| A594<br>gt anti-rat | Thermo Fisher | A11007<br>AB_10561522 | 1:400 |
| A546<br>gt anti-ms | Thermo Fisher | A11003<br>AB_2534071 | 1:400 |
| A488<br>gt anti-rb | Thermo Fisher | A11008<br>AB_143165 | 1:400 |
| A488<br>gt anti-chk | Thermo Fisher | A11039<br>AB_2534096 | 1:400 |
| anti-P2X7-A647 | Nolte lab | - | 1:100 |
| anti-Ly6G-A700 | Biolegend | 127622 | 1:100 |
| anti-CD3eBv650 | Biolegend | 100229 | 1:100 |
| anti-CD8a-Bv711 | Biolegend | 100747 | 1:100 |
| anti-CD4-Bv605 | Biolegend | 100547 | 1:100 |
| anti-gdTCR-Bv421 | Biolegend | 118119 | 1:100 |
| anti-CD11b-APC-Cy7 | Biolegend | 101225 | 1:100 |
| anti-CD11c-PE-Dazzle | Biolegend | 117347 | 1:100 |
| anti-CD64-PE-Cy7 | Biolegend | 139313 | 1:100 |

## References

Antonioli, L., Blandizzi, C., and Giron, M.C. (2015). Enteric purinergic signaling: Shaping the “brain in the gut”. Neuropharmacology 95, 477–478. 10.1016/j.neuropharm.2015.04.021.

Antonioli, L., Blandizzi, C., Pacher, P., and Hasko, G. (2019). The Purinergic System as a Pharmacological Target for the Treatment of Immune-Mediated Inflammatory Diseases. Pharmacol Rev 71, 345–382. 10.1124/pr.117.014878.

Antonioli, L., Giron, M.C., Colucci, R., Pellegrini, C., Sacco, D., Caputi, V., Orso, G., Tuccori, M., Scarpignato, C., Blandizzi, C., and Fornai, M. (2014). Involvement of the P2X7 purinergic receptor in colonic motor dysfunction associated with bowel inflammation in rats. PLoS One 9, e116253. 10.1371/journal.pone.0116253.

Baumgart, D.C., and Carding, S.R. (2007). Inflammatory bowel disease: cause and immunobiology. Lancet 369, 1627–1640. 10.1016/S0140-6736(07)60750-8.

Bhave, S., Gade, A., Kang, M., Hauser, K.F., Dewey, W.L., and Akbarali, H.I. (2017). Connexin-purinergic signaling in enteric glia mediates the prolonged effect of morphine on constipation. FASEB J 31, 2649–2660. 10.1096/fj.201601068R.

Boesmans, W., Cirillo, C., Van den Abbeel, V., Van den Haute, C., Depoortere, I., Tack, J., and Vanden Berghe, P. (2013). Neurotransmitters involved in fast excitatory neurotransmission directly activate enteric glial cells. Neurogastroenterol Motil 25, e151–160. 10.1111/nmo.12065.

Boesmans, W., Lasrado, R., Vanden Berghe, P., and Pachnis, V. (2015). Heterogeneity and phenotypic plasticity of glial cells in the mammalian enteric nervous system. Glia 63, 229–241. 10.1002/glia.22746.

Brown, I.A., McClain, J.L., Watson, R.E., Patel, B.A., and Gulbransen, B.D. (2016). Enteric glia mediate neuron death in colitis through purinergic pathways that require connexin-43 and nitric oxide. Cell Mol Gastroenterol Hepatol 2, 77–91. 10.1016/j.jcmgh.2015.08.007.

Burnstock, G. (2016). Purinergic Signalling in the Gut. Adv Exp Med Biol 891, 91–112. 10.1007/978-3-319-27592-5_10.

Cesaro, A., Brest, P., Hofman, V., Hebuterne, X., Wildman, S., Ferrua, B., Marchetti, S., Doglio, A., Vouret-Craviari, V., Galland, F., et al. (2010). Amplification loop of the inflammatory process is induced by P2X7R activation in intestinal epithelial cells in response to neutrophil transepithelial migration. Am J Physiol Gastrointest Liver Physiol 299, G32–42. 10.1152/ajpgi.00282.2009.

da Silva, M.V., Marosti, A.R., Mendes, C.E., Palombit, K., and Castelucci, P. (2017). Submucosal neurons and enteric glial cells expressing the P2X7 receptor in rat experimental colitis. Acta Histochem 119, 481–494. 10.1016/j.acthis.2017.05.001.

Desmet, A.S., Cirillo, C., and Vanden Berghe, P. (2014). Distinct subcellular localization of the neuronal marker HuC/D reveals hypoxia-induced damage in enteric neurons. Neurogastroenterol Motil 26, 1131–1143. 10.1111/nmo.12371.

Di Virgilio, F., Dal Ben, D., Sarti, A.C., Giuliani, A.L., and Falzoni, S. (2017). The P2X7 Receptor in Infection and Inflammation. Immunity 47, 15–31. 10.1016/j.immuni.2017.06.020.

Di Virgilio, F., Sarti, A.C., and Grassi, F. (2018). Modulation of innate and adaptive immunity by P2X ion channels. Curr Opin Immunol 52, 51–59. 10.1016/j.coi.2018.03.026.

Diezmos, E.F., Bertrand, P.P., and Liu, L. (2016). Purinergic Signaling in Gut Inflammation: The Role of Connexins and Pannexins. Front Neurosci 10, 311. 10.3389/fnins.2016.00311.

Diezmos, E.F., Markus, I., Perera, D.S., Gan, S., Zhang, L., Sandow, S.L., Bertrand, P.P., and Liu, L. (2018). Blockade of Pannexin-1 Channels and Purinergic P2X7 Receptors Shows Protective Effects Against Cytokines-Induced Colitis of Human Colonic Mucosa. Front Pharmacol 9, 865. 10.3389/fphar.2018.00865.

Duveau, A., Bertin, E., and Boue-Grabot, E. (2020). Implication of Neuronal Versus Microglial P2X4 Receptors in Central Nervous System Disorders. Neurosci Bull 36, 1327–1343. 10.1007/s12264-020-00570-y.

Eser, A., Colombel, J.F., Rutgeerts, P., Vermeire, S., Vogelsang, H., Braddock, M., Persson, T., and Reinisch, W. (2015). Safety and Efficacy of an Oral Inhibitor of the Purinergic Receptor P2X7 in Adult Patients with Moderately to Severely Active Crohn’s Disease: A Randomized Placebo-controlled, Double-blind, Phase IIa Study. Inflamm Bowel Dis 21, 2247–2253. 10.1097/MIB.0000000000000514.

Evangelinellis, M.M., Souza, R.F., Mendes, C.E., and Castelucci, P. (2021). Effects of a P2X7 receptor antagonist on myenteric neurons in the distal colon of an experimental rat model of ulcerative colitis. Histochem Cell Biol. 10.1007/s00418-021-02039-z.

Figliuolo, V.R., Savio, L.E.B., Safya, H., Nanini, H., Bernardazzi, C., Abalo, A., de Souza, H.S.P., Kanellopoulos, J., Bobe, P., Coutinho, C., and Coutinho-Silva, R. (2017). P2X7 receptor promotes intestinal inflammation in chemically induced colitis and triggers death of mucosal regulatory T cells. Biochim Biophys Acta Mol Basis Dis 1863, 1183–1194. 10.1016/j.bbadis.2017.03.004.

Gabanyi, I., Muller, P.A., Feighery, L., Oliveira, T.Y., Costa-Pinto, F.A., and Mucida, D. (2016). Neuro-immune Interactions Drive Tissue Programming in Intestinal Macrophages. Cell 164, 378–391. 10.1016/j.cell.2015.12.023.

Galligan, J.J., and North, R.A. (2004). Pharmacology and function of nicotinic acetylcholine and P2X receptors in the enteric nervous system. Neurogastroenterol Motil 16 Suppl 1, 64–70. 10.1111/j.1743-3150.2004.00478.x.

Ginhoux, F., and Prinz, M. (2015). Origin of microglia: current concepts and past controversies. Cold Spring Harb Perspect Biol 7, a020537. 10.1101/cshperspect.a020537.

Gomes, P., Chevalier, J., Boesmans, W., Roosen, L., van den Abbeel, V., Neunlist, M., Tack, J., and Vanden Berghe, P. (2009). ATP-dependent paracrine communication between enteric neurons and glia in a primary cell culture derived from embryonic mice. Neurogastroenterol Motil 21, 870–e862. 10.1111/j.1365-2982.2009.01302.x.

Grubisic, V., McClain, J.L., Fried, D.E., Grants, I., Rajasekhar, P., Csizmadia, E., Ajijola, O.A., Watson, R.E., Poole, D.P., Robson, S.C., et al. (2020). Enteric Glia Modulate Macrophage Phenotype and Visceral Sensitivity following Inflammation. Cell Rep 32, 108100. 10.1016/j.celrep.2020.108100.

Gulbransen, B.D., Bashashati, M., Hirota, S.A., Gui, X., Roberts, J.A., MacDonald, J.A., Muruve, D.A., McKay, D.M., Beck, P.L., Mawe, G.M., et al. (2012). Activation of neuronal P2X7 receptor-pannexin-1 mediates death of enteric neurons during colitis. Nat Med 18, 600–604. 10.1038/nm.2679.

Gulbransen, B.D., and Sharkey, K.A. (2009). Purinergic neuron-to-glia signaling in the enteric nervous system. Gastroenterology 136, 1349–1358. 10.1053/j.gastro.2008.12.058.

Gulbransen, B.D., and Sharkey, K.A. (2012). Novel functional roles for enteric glia in the gastrointestinal tract. Nat Rev Gastroenterol Hepatol 9, 625–632. 10.1038/nrgastro.2012.138.

Hofman, P., Cherfils-Vicini, J., Bazin, M., Ilie, M., Juhel, T., Hebuterne, X., Gilson, E., Schmid-Alliana, A., Boyer, O., Adriouch, S., and Vouret-Craviari, V. (2015). Genetic and pharmacological inactivation of the purinergic P2RX7 receptor dampens inflammation but increases tumor incidence in a mouse model of colitis-associated cancer. Cancer Res 75, 835–845. 10.1158/0008-5472.CAN-14-1778.

Kaczmarek-Hajek, K., Lorinczi, E., Hausmann, R., and Nicke, A. (2012). Molecular and functional properties of P2X receptors--recent progress and persisting challenges. Purinergic Signal 8, 375–417. 10.1007/s11302-012-9314-7.

Kaczmarek-Hajek, K., Zhang, J., Kopp, R., Grosche, A., Rissiek, B., Saul, A., Bruzzone, S., Engel, T., Jooss, T., Krautloher, A., et al. (2018). Re-evaluation of neuronal P2X7 expression using novel mouse models and a P2X7-specific nanobody. Elife 7. 10.7554/eLife.36217.

Kopp, R., Krautloher, A., Ramirez-Fernandez, A., and Nicke, A. (2019). P2X7 Interactions and Signaling - Making Head or Tail of It. Front Mol Neurosci 12, 183. 10.3389/fnmol.2019.00183.

Kurashima, Y., Amiya, T., Nochi, T., Fujisawa, K., Haraguchi, T., Iba, H., Tsutsui, H., Sato, S., Nakajima, S., Iijima, H., et al. (2012). Extracellular ATP mediates mast cell-dependent intestinal inflammation through P2X7 purinoceptors. Nat Commun 3, 1034. 10.1038/ncomms2023.

Linan-Rico, A., Wunderlich, J.E., Enneking, J.T., Tso, D.R., Grants, I., Williams, K.C., Otey, A., Michel, K., Schemann, M., Needleman, B., et al. (2015). Neuropharmacology of purinergic receptors in human submucous plexus: Involvement of P2X(1), P2X(2), P2X(3) channels, P2Y and A(3) metabotropic receptors in neurotransmission. Neuropharmacology 95, 83–99. 10.1016/j.neuropharm.2015.02.014.

Matsukawa, T., Izawa, K., Isobe, M., Takahashi, M., Maehara, A., Yamanishi, Y., Kaitani, A., Okumura, K., Teshima, T., Kitamura, T., and Kitaura, J. (2016). Ceramide-CD300f binding suppresses experimental colitis by inhibiting ATP-mediated mast cell activation. Gut 65, 777–787. 10.1136/gutjnl-2014-308900.

Mendes, C.E., Palombit, K., Tavares-de-Lima, W., and Castelucci, P. (2019). Enteric glial cells immunoreactive for P2X7 receptor are affected in the ileum following ischemia and reperfusion. Acta Histochem 121, 665–679. 10.1016/j.acthis.2019.06.001.

Morales-Soto, W., and Gulbransen, B.D. (2019). Enteric Glia: A New Player in Abdominal Pain. Cell Mol Gastroenterol Hepatol 7, 433–445. 10.1016/j.jcmgh.2018.11.005.

Muller, P.A., Koscso, B., Rajani, G.M., Stevanovic, K., Berres, M.L., Hashimoto, D., Mortha, A., Leboeuf, M., Li, X.M., Mucida, D., et al. (2014). Crosstalk between muscularis macrophages and enteric neurons regulates gastrointestinal motility. Cell 158, 300–313. 10.1016/j.cell.2014.04.050.

Na, Y.R., Stakenborg, M., Seok, S.H., and Matteoli, G. (2019). Macrophages in intestinal inflammation and resolution: a potential therapeutic target in IBD. Nat Rev Gastroenterol Hepatol 16, 531–543. 10.1038/s41575-019-0172-4.

Niesler, B., Kuerten, S., Demir, I.E., and Schafer, K.H. (2021). Disorders of the enteric nervous system - a holistic view. Nat Rev Gastroenterol Hepatol. 10.1038/s41575-020-00385-2.

Nieto-Pescador, M.G., Guerrero-Alba, R., Valdez-Morales, E., Espinosa-Luna, R., Jimenez-Vargas, N., Linan-Rico Andromeda, A., Ramos-Lomas, T.L., Diaz-Hernandez Veronica, V., Montano, L.M., and Barajas-Lopez, C. (2013). P2X4 subunits are part of P2X native channels in murine myenteric neurons. Eur J Pharmacol 709, 93–102. 10.1016/j.ejphar.2013.03.045.

Ohbori, K., Fujiwara, M., Ohishi, A., Nishida, K., Uozumi, Y., and Nagasawa, K. (2017). Prophylactic Oral Administration of Magnesium Ameliorates Dextran Sulfate Sodium-Induced Colitis in Mice through a Decrease of Colonic Accumulation of P2X7 Receptor-Expressing Mast Cells. Biol Pharm Bull 40, 1071–1077. 10.1248/bpb.b17-00143.

Orholm, M., Munkholm, P., Langholz, E., Nielsen, O.H., Sorensen, T.I., and Binder, V. (1991). Familial occurrence of inflammatory bowel disease. N Engl J Med 324, 84–88. 10.1056/NEJM199101103240203.

Palombit, K., Mendes, C.E., Tavares-de-Lima, W., Barreto-Chaves, M.L., and Castelucci, P. (2019). Blockage of the P2X7 Receptor Attenuates Harmful Changes Produced by Ischemia and Reperfusion in the Myenteric Plexus. Dig Dis Sci 64, 1815–1829. 10.1007/s10620-019-05496-8.

Phillips, R.J., and Powley, T.L. (2012). Macrophages associated with the intrinsic and extrinsic autonomic innervation of the rat gastrointestinal tract. Auton Neurosci 169, 12–27. 10.1016/j.autneu.2012.02.004.

Proietti, M., Cornacchione, V., Rezzonico Jost, T., Romagnani, A., Faliti, C.E., Perruzza, L., Rigoni, R., Radaelli, E., Caprioli, F., Preziuso, S., et al. (2014). ATP-gated ionotropic P2X7 receptor controls follicular T helper cell numbers in Peyer’s patches to promote host-microbiota mutualism. Immunity 41, 789–801. 10.1016/j.immuni.2014.10.010.

Ren, J., Bian, X., DeVries, M., Schnegelsberg, B., Cockayne, D.A., Ford, A.P., and Galligan, J.J. (2003). P2X2 subunits contribute to fast synaptic excitation in myenteric neurons of the mouse small intestine. J Physiol 552, 809–821. 10.1113/jphysiol.2003.047944.

Rissiek, B., Haag, F., Boyer, O., Koch-Nolte, F., and Adriouch, S. (2015). P2X7 on Mouse T Cells: One Channel, Many Functions. Front Immunol 6, 204. 10.3389/fimmu.2015.00204.

Rumney, R.M.H., and Gorecki, D.C. (2020). Knockout and Knock-in Mouse Models to Study Purinergic Signaling. Methods Mol Biol 2041, 17–43. 10.1007/978-1-4939-9717-6_2.

Sanchez-Munoz, F., Dominguez-Lopez, A., and Yamamoto-Furusho, J.K. (2008). Role of cytokines in inflammatory bowel disease. World J Gastroenterol 14, 4280–4288. 10.3748/wjg.14.4280.

Schenk, U., Frascoli, M., Proietti, M., Geffers, R., Traggiai, E., Buer, J., Ricordi, C., Westendorf, A.M., and Grassi, F. (2011). ATP inhibits the generation and function of regulatory T cells through the activation of purinergic P2X receptors. Sci Signal 4, ra12. 10.1126/scisignal.2001270.

Sharkey, K.A., Beck, P.L., and McKay, D.M. (2018). Neuroimmunophysiology of the gut: advances and emerging concepts focusing on the epithelium. Nat Rev Gastroenterol Hepatol 15, 765–784. 10.1038/s41575-018-0051-4.

Spear, E.T., and Mawe, G.M. (2019). Enteric neuroplasticity and dysmotility in inflammatory disease: key players and possible therapeutic targets. Am J Physiol Gastrointest Liver Physiol 317, G853–G861. 10.1152/ajpgi.00206.2019.

Van Crombruggen, K., Van Nassauw, L., Timmermans, J.P., and Lefebvre, R.A. (2007). Inhibitory purinergic P2 receptor characterisation in rat distal colon. Neuropharmacology 53, 257–271. 10.1016/j.neuropharm.2007.05.010.

Vanderwinden, J.M., Timmermans, J.P., and Schiffmann, S.N. (2003). Glial cells, but not interstitial cells, express P2X7, an ionotropic purinergic receptor, in rat gastrointestinal musculature. Cell Tissue Res 312, 149–154. 10.1007/s00441-003-0716-2.

Verheijden, S., De Schepper, S., and Boeckxstaens, G.E. (2015). Neuron-macrophage crosstalk in the intestine: a “microglia” perspective. Front Cell Neurosci 9, 403. 10.3389/fncel.2015.00403.

Viola, M.F., and Boeckxstaens, G. (2020). Intestinal resident macrophages: Multitaskers of the gut. Neurogastroenterol Motil 32, e13843. 10.1111/nmo.13843.

Vuerich, M., Mukherjee, S., Robson, S.C., and Longhi, M.S. (2020). Control of Gut Inflammation by Modulation of Purinergic Signaling. Front Immunol 11, 1882. 10.3389/fimmu.2020.01882.

Xavier, R.J., and Podolsky, D.K. (2007). Unravelling the pathogenesis of inflammatory bowel disease. Nature 448, 427–434. 10.1038/nature06005.

