## Supplement for "Macrophages and glia are the dominant P2X7-expressing cell types in the gut nervous system – no evidence for a role of neuronal P2X7 receptors in colitis"

**Suppl. Figure 1: Identification of two types of macrophages by triple staining of LMMP preparations for P2X7-EGFP, the macrophage marker CD68, and Iba1.** Ramified macrophages associated with ganglia were stained with all three antibodies. Cd68- and P2X7-EGFP-positive and Iba1-gegative macrophages (indicated by arrows in the right image) show extended morphology and parallel orientation.

**Suppl. Figure 2: P2X7 is not detected in CD117 and Ano1- positive cells of Cajal.** LMMP preparations from P2X7-EGFP transgenic and wt mice were stained for P2X7 or EGFP and antibodies against markers for cells of Cajal (CD117 and Ano1) as indicated.

**Suppl. Figure 3: The P2X7 nanobody shows specific P2X7 staining that is mirrored in P2X7-EGFP transgenic mice.** LMMP preparations from P2X7-EGFP, wt, and *P2rx7*$^{-/-}$ were stained with the P2X7 nanobody and antibodies against the neuronal marker HuC/D.

**Suppl. Figure 4: P2X4 co-localizes with the lysosome/macrophage marker CD68 in macrophages of the myenteric plexus.**

Supplementary Fig. 1

A

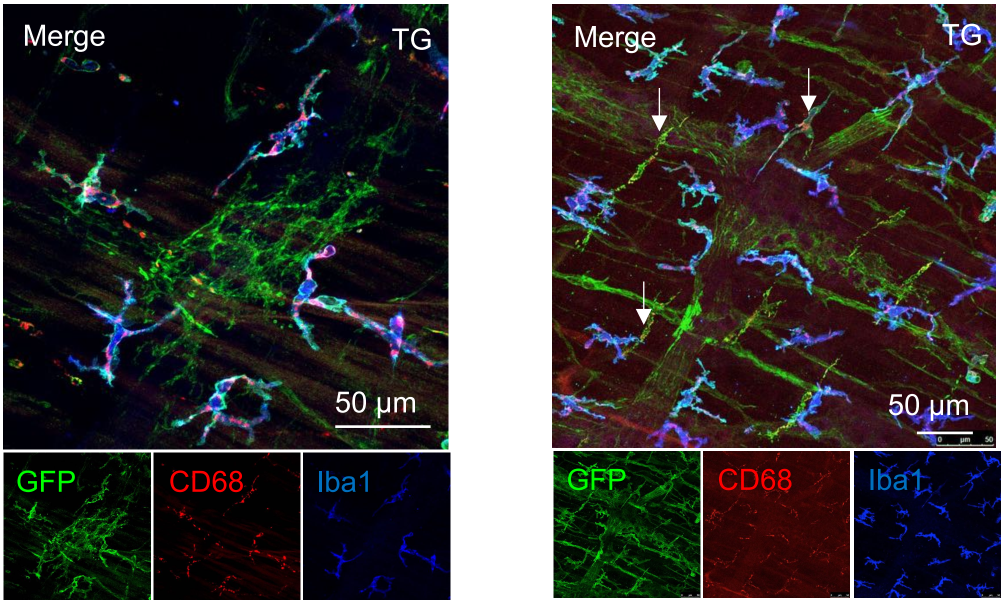

Supplementary Fig. 2

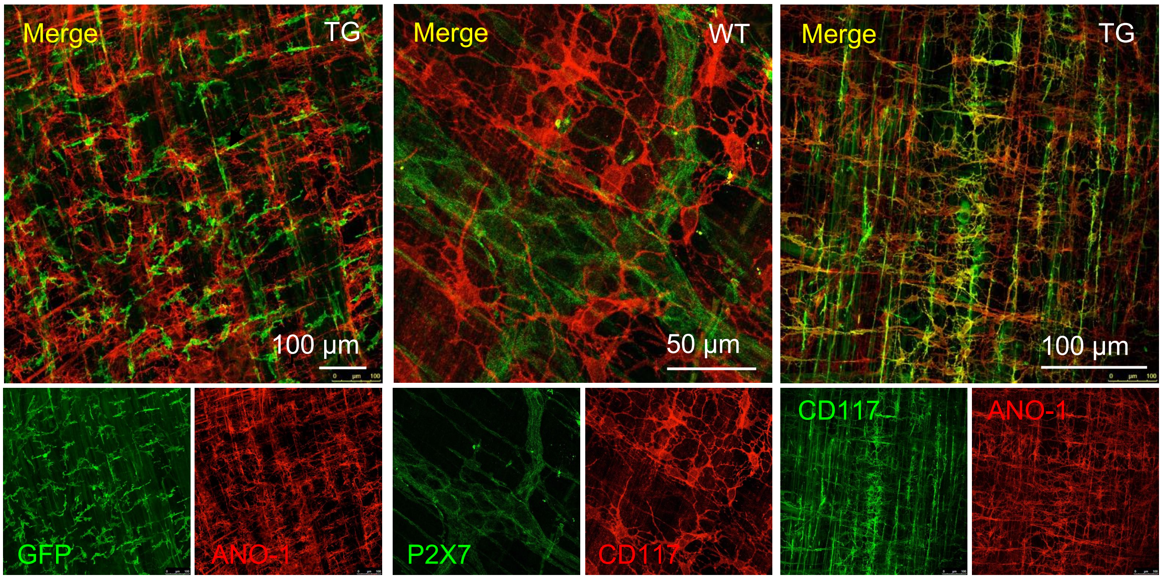

Supplementary Fig. 3

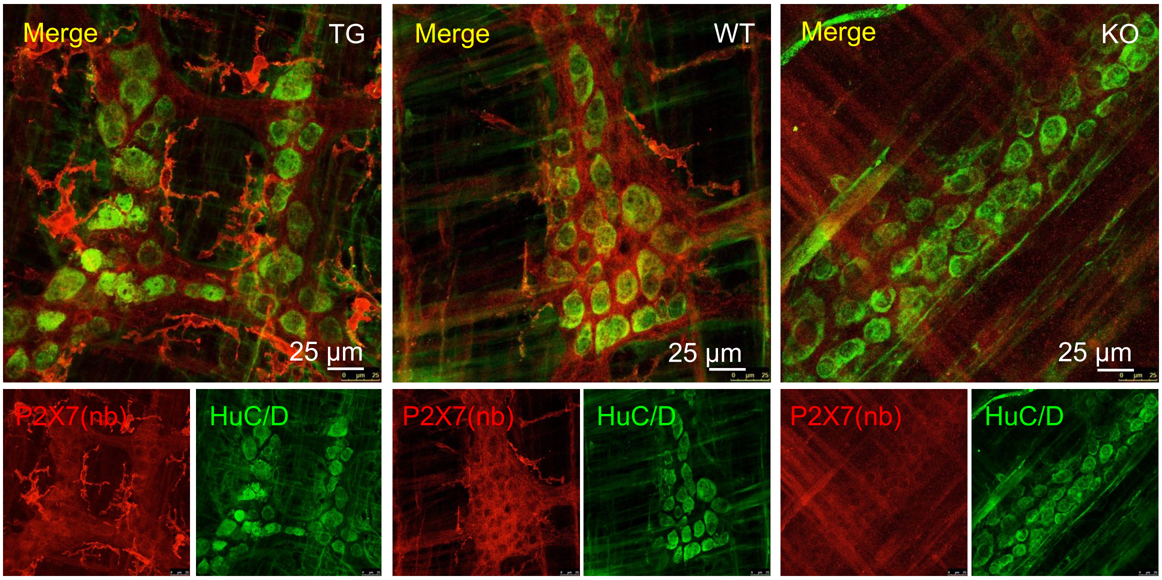

Supplementary Fig. 4

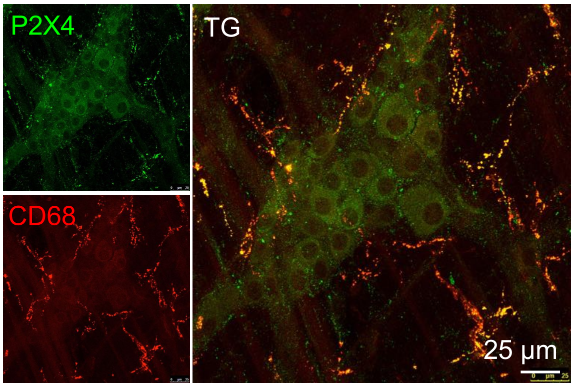
